# Gene Co-expression Connectivity Predicts Gene Targets Underlying High Ionic Liquid Tolerance in *Yarrowia lipolytica*

**DOI:** 10.1101/2022.03.17.484694

**Authors:** Caleb Walker, Seunghyun Ryu, Sergio Garcia, David Dooley, Brian Mendoza, Cong T. Trinh

## Abstract

Microbial tolerance to organic solvents such as ionic liquids (ILs) is a robust phenotype beneficial for novel biotransformation. While most microbes become inhibited in 1%-5% (v/v) IL (e.g., 1-ethyl-3-methylimidazolium acetate), we engineered a robust *Yarrowia lipolytica* (YlCW001) that tolerates a record high of 18% (v/v) IL via adaptive laboratory evolution. Yet, genotypes conferring high IL tolerance in YlCW001 remain to be discovered. Using dynamic RNA-Seq data, we shed light on the underlying cellular processes that *Y. lipolytica* responds to IL. We introduced Gene Co-expression Connectivity (GeCCo) to discover genotypes conferring desirable phenotypes that might not be found by the conventional differential expression (DE) approaches. GeCCo selects genes based on their number of co-expressed genes in a sub-network of upregulated genes by the target phenotype. We experimentally validated GeCCo by reversely engineering high IL-tolerant phenotype in wildtype *Y. lipolytica*. We found that gene targets selected by both DE and GeCCo exhibited the best statistical chance at increasing IL tolerance when individually overexpressed. Remarkably, the best combination of dual-overexpressed genes were genes selected by GeCCo alone. This non-intuitive combination of genes, BRN1 and OYE2, are involved in guiding/regulating mitotic cell division, chromatin segregation/condensation, microtubule and cytoskeletal organization, and Golgi vesicle transport.

## BACKGROUND

Cellular robustness such as solvent tolerance is an important phenotype in natural and engineered biological systems. Biocatalysis in organic solvents enables novel biotransformation with unique strategies for substrate solubilization (1), enhanced enzymatic activity (2), and product recovery (3). Ionic liquids (ILs) have recently emerged as green organic solvents to replace the conventional ones in bioprocessing (4). These solvents can provide a novel reaction medium for biotransformation with superior performance due to their ability to dissolve a broader range of compounds and their adjustable properties for enzyme stabilization and activation (e.g., lipases, alcohol dehydrogenases, proteases, and oxidoreductases) (5). Some ILs such as 1-ethyl-3-methylimidazolium acetate ([EMIM][OAc]) are promising for the cellulosic biomass pretreatment technology that effectively reduces recalcitrance, achieving efficient dissolution of biomass and permitting enzymatic accessibility to the sugar polymers (i.e., cellulose and hemicellulose) for saccharification (6–8). Fermentative sugars released from IL-pretreated biomass could be assimilated by microbial cell factories if not for the toxicity that ILs impose on microorganisms (9, 10). For these reasons, microbial high solvent tolerance is a desirable phenotype for conversion of renewable cellulosic biomass to replace petroleum-derived chemicals and fuels.

ILs are toxic to most industrial workhorse microorganisms (e.g., *Escherichia coli, Saccharomyces cerevisiae*). Cell growth is inhibited even at low IL concentrations, for example, 1-5% (v/v) [EMIM][OAc] (11, 12). These ILs disrupt cell membranes and intracellular processes (13–15). Efflux pumps have been reported to improve IL tolerance in microorganisms (9, 16–19). Currently, knowledge of novel genotypes responsible for IL tolerance in microorganisms is still very limited. Remarkably, screens for IL-tolerant microorganisms identified the GRAS (generally-regarded-as-safe) oleaginous yeast, *Yarrowia lipolytica*, as one of the top performers in the benchmark [EMIM][OAc] (20). Wildtype *Y. lipolytica* could achieve 92% of the theoretical yield of alpha ketoglutaric acid (KGA) in 10% (v/v) [EMIM][OAc] during simultaneous saccharification and fermentation of IL-pretreated cellulose (21). Due to its robustness and broad industrial use, *Y. lipolytica* is a promising model organism to study IL robustness.

To understand robustness of *Y. lipolytica* in ILs, adaptive laboratory evolution (ALE) was performed to generate a platform strain YlCW001 with enhanced activity in high concentrations of 18% (v/v) [EMIM][OAc] (22). Remarkably, the mutant also outperformed the wildtype *Y. lipolytica* in all imidazolium-based ILs tested. Physiological, metabolic, and genetic characterization elucidated that sterols are critical for IL tolerance in *Y. lipolytica* by strengthening cell membrane (22). Despite the discovery of beneficial role of sterols, other novel genotypes that confer IL tolerance in *Y. lipolytica* still remain to be discovered. Through genome-wide analysis of the wildtype and evolved mutant *Y. lipolytica* exhibiting different degrees of IL tolerance, it is promising to elucidate the underlying beneficial genotypes. The key challenge is however to identify and validate gene targets and their combinations among a large set of gene candidates in response to high IL exposure. Since IL tolerance is a complex phenotype, the most differential expression genes identified by OMICS data (e.g., RNAseq) alone might not necessarily be the prime candidates conferring IL tolerance.

In this study, we seek to identify key genotype(s) conferring IL tolerance in *Y. lipolytica*. Using dynamic transcriptomic data of the wildtype and evolved *Y. lipolytica* strains growing with and without IL, we shed light on the underlying cellular processes that *Y. lipolytica* responds to IL. We further developed Gene Co-expression Connectivity (GeCCo) as a metric to identify a small subset of promising gene targets conferring IL tolerance among a large set of potential gene candidates. Specifically, GeCCo selects the most connected genes from a co-expression network of upregulated genes. By performing single and dual gene overexpression validation, we demonstrated that GeCCo could reveal key genotype(s) behind complex phenotype(s) that were reverse engineered to achieve high solvent-tolerance in *Y. lipolytica*.

## RESULTS

### Modified genotype of the IL-tolerant evolved strain YlCW001

To connect genetic mutations with enhanced IL tolerance, we re-sequenced the evolved strain YlCW001 (MT) and analyzed the mutations. Across the entire genome, we detected a total of 648 variants but only 40 genes contained variants that caused an amino acid change (Figure 1A, Supplementary File S2). These 40 mutated genes (MGs) are randomly distributed across the 6 chromosomes of *Y. lipolytica* and held a total of 68 mutations (i.e., variants) including 35 single nucleotide variants (SNV), 12 deletions, 11 multiple nucleotide variants (MNV), 8 insertions and 2 replacements (Figure 1B). The most MGs were in chromosome A (13 MGs) followed by chromosome E (9 MGs), chromosome F (7 MGs), chromosome C (6 MGs), chromosomes B and D (each with 3 MGs) (Figure 1A). By performing gene ontology (GO) associations for the 40 MGs using cluego, IMG, and blastP against *S. cerevisiae*, we were able to assign the functional annotations for 68% of the MGs (Figure 1C). Kinases represent the most abundant GO terms containing 3 MGs, followed by transporters, transcription factors, mRNA processing, regulation, nuclear pore complex and membrane components (2 MGs each, see Supplementary File S2 for gene IDs).

**Figure 1.**
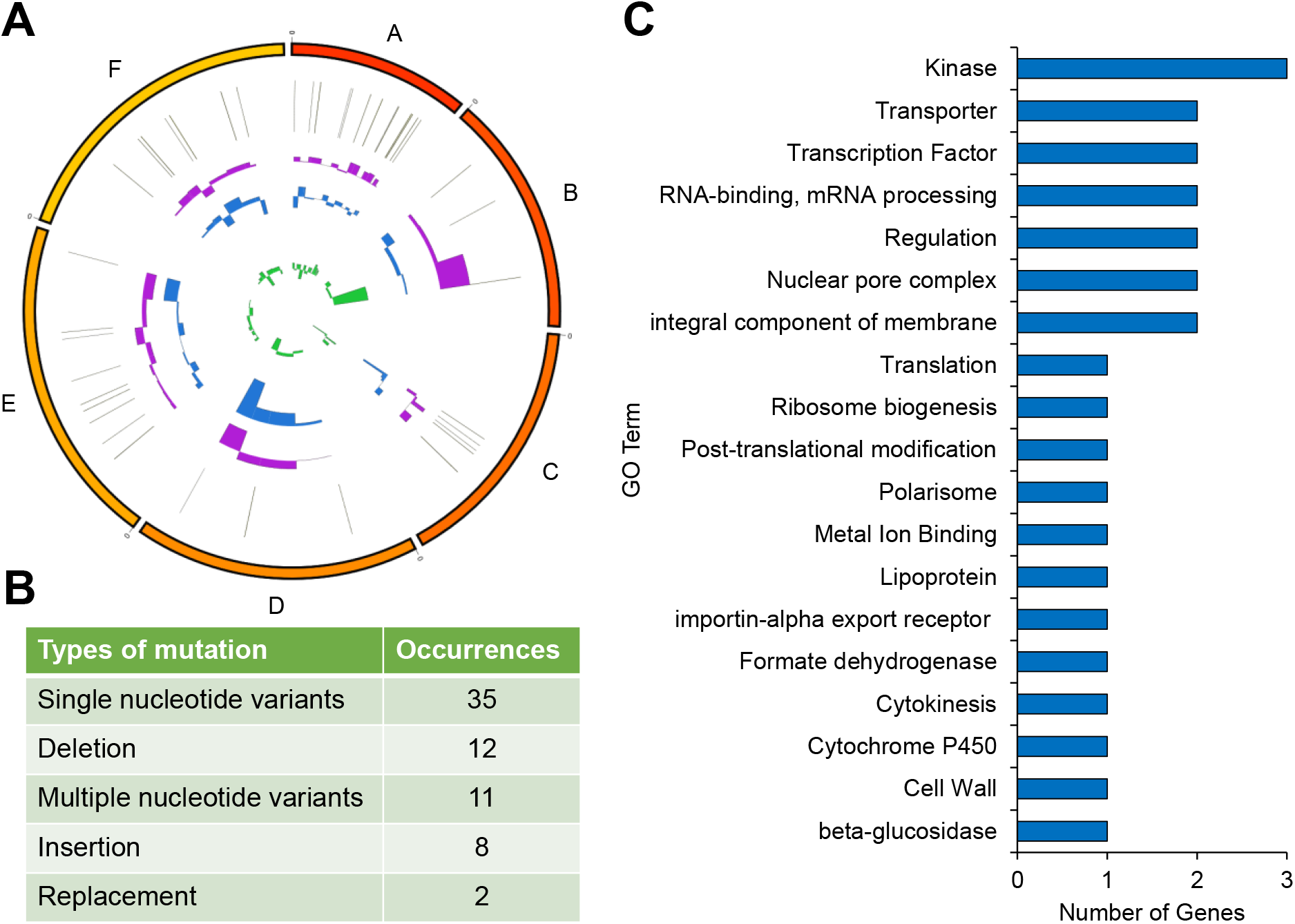
Mutated genes of IL-tolerant mutant *Y. lipolyti*ca (MT, YlCW001) which include 68 variants across 40 genes that change amino acid residues. **(A)** Chromosomal location of mutated genes depicting log_2_ fold changes at the mid-exponential RNA-seq sample between the MT in 8% [EMIM][OAc] versus WT in 0% IL (purple), WT in 8% IL (blue) and MT in 0% IL (green). **(B)** Types of variants in the mutated genes. **(C)** Gene ontology of the mutated genes.

### Basal gene expression differences between the WT and MT strains without IL exposure

We first examined the basal changes in gene expression between the WT and MT strains growing in medium without IL (Figure 2A). Temporal RNAseq data for both the WT and MT strains were collected at the early and mid-exponential growth phases and subjected to the gene classification analysis (Supplementary File S3, data set i). We identified 470 “upregulated” genes and 88 “increasing” genes in the MT strain relative to the WT strain cultured without IL (Figure 2E). Here, upregulated genes are overexpressed at both early and mid-exponential phases while increasing genes are overexpressed at mid-exponential phase but do not exhibit significant change at early-exponential phase (see Materials and Methods). These genes with greater expression in the MT strain were annotated for mitochondrion, nucleus, chromosome, membrane protein complex, microbody, respirasome and transmembrane signal receptors (Figure 2I). For the WT strain, we found in 329 upregulated genes and 167 increasing genes relative to the MT strain cultured without IL. These genes were involved in cell periphery, cell cortex, cell wall organization/biogenesis, endoplasmic reticulum, and the Golgi apparatus (Figure 2I). Interestingly, 5 of the MGs were differentially expressed between the MT and WT cultured without IL. The MT upregulated 3 of these MGs including an RNA metabolic protein (REH1, YALI0B08734g), an RNS splicing factor (WHI1, YALI0F12375g), and an uncharacterized protein (YALI0C13002g). Meanwhile, the MT downregulated 2 of these MGs, an expansin-like protein (YALI0E17941g) and a non-receptor serine/threonine protein kinase (SKY1, YALI0A18590g).

**Figure 2.**
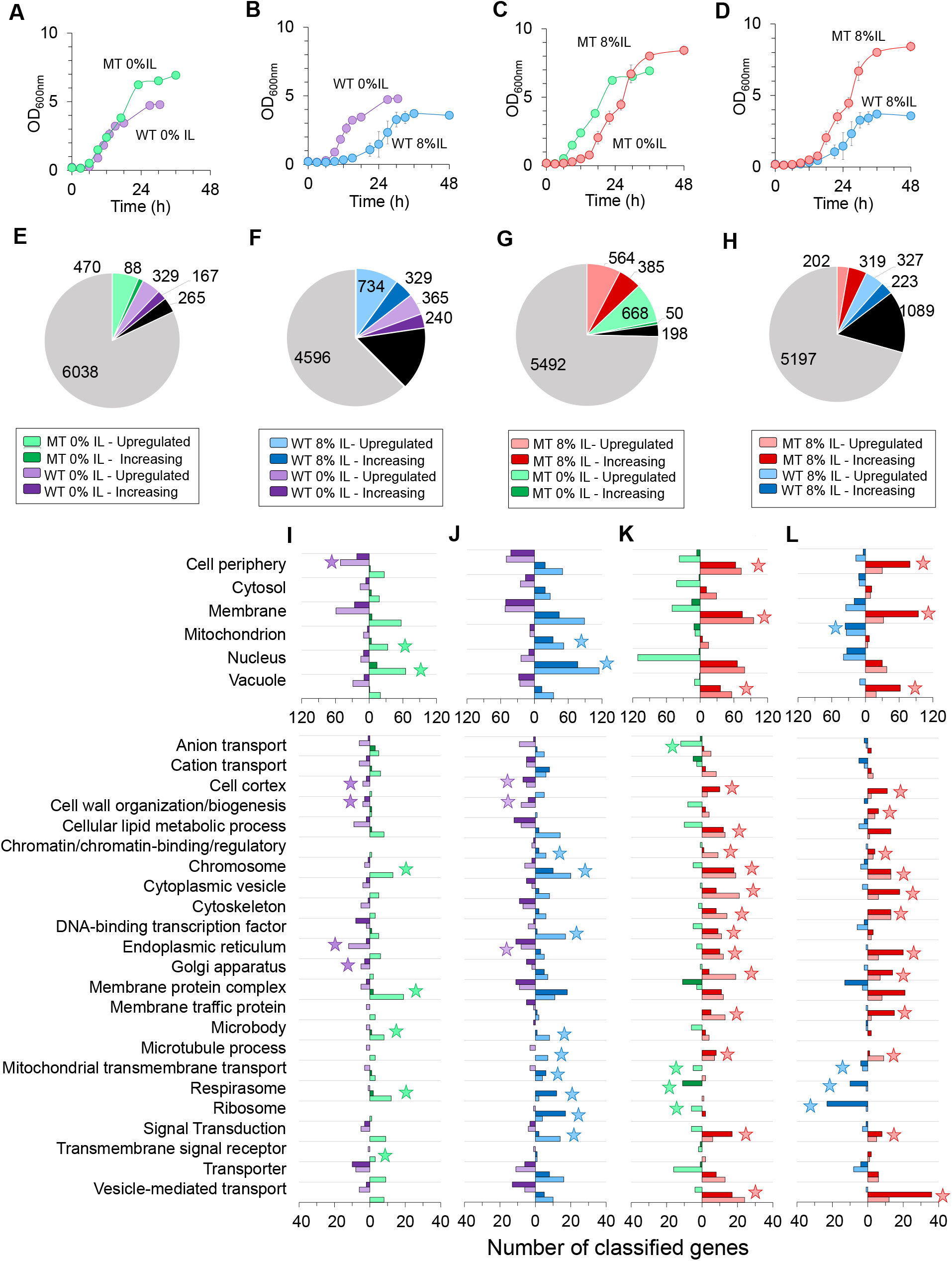
Growth profiles of (**A**) MT and WT in 0% IL, (**B**) WT in 0% IL and WT in 8% (v/v) IL, (**C**) MT in 0% IL and MT in 8% (v/v) IL, and (**D**) Growth of WT and MT in 8% (v/v) IL. (**E-H**) Number and distribution of classified genes for (**E**) MT 0% IL vs WT 0% IL, (**F**) WT 8% IL vs WT 0% IL, (**G**) MT 8% IL vs MT 0% IL, and (H) MT 8% IL vs WT 8% IL. In these pie charts, black and gray pies are referred to “changed regulation” and “no change” genes, respectively. (**I-L**) Classified gene ontology associations for (**I**) basal differences in gene expression between the MT and WT in 0% IL, (**J**) IL-responsive gene expression of the WT strain, (**K**) IL-responsive gene expression of the MT strain, and (**L**) enhanced IL-responsive gene expression of the MT strain versus the WT strain in 8% IL. Stars represent ontology terms where the sum of upregulated and increasing genes in a biological condition is at least twice greater than the other biological condition.

### IL-responsive gene expression

#### Wildtype response to IL

Next, we aimed to understand gene expression changes for the WT strain exposed to IL (Figure 2B) (Supplementary file S3, data set ii). This RNA-seq pairwise set exhibited the greatest perturbation of gene expression relative to all other data sets with only ∼62% of genes classified as no change (Figure 2F). We found 734 upregulated genes and 329 increasing genes for the WT growing in 8% (v/v) IL. These IL-induced genes were enriched for mitochondrion (including mitochondrial transmembrane transport and respirasome), genetic processing (i.e., nucleus, chromosome, chromatin-binding/regulation, DNA-binding transcription factors, ribosome, and signal transduction), microtubule processing, and microbody (Figure 2J). In contrast, we identified 365 upregulated genes and 240 increasing genes for the WT cultured without IL. However, only 3 annotated processes were enriched more than 2-fold which included cell cortex, cell wall organization/biogenesis and the endoplasmic reticulum (Figure 2J).

#### Mutant response to IL

Likewise, we compared gene expression changes for the MT strain exposed to IL (Figure 2C) (Supplementary File S3, data set iii). The gene classification analysis identified 564 upregulated genes and 385 increasing genes for the MT strain growing in 8% (v/v) IL relative to the MT strain without IL (Figure 2G). These IL-induced genes for the MT strain were enriched for lipid processes (i.e., membrane, cellular lipid metabolic process, membrane traffic protein, Golgi apparatus, endoplasmic reticulum), cell periphery (including cell cortex, cytoskeleton and microtubule process), intracellular vesicles (i.e., vacuole, cytoplasmic vesicle, vesicle-mediated transport), and genetic processes (i.e., chromosome, chromatin/chromatin-binding/regulator, DNA-binding transcription factor, signal transduction) (Figure 2K). The MT strain cultured without IL exhibited 668 upregulated genes and only 50 increasing genes. These gene sets were annotated for anion and mitochondrial transmembrane transport, respirasome and the ribosome (Figure 2K). We also identified 9 of the MGs among the classified genes. Of these, 5 were upregulated in IL including an actin cytoskeletal protein (YALI0A18381g), a non-receptor serine/threonine protein kinase (CBK1, YALI0B04268g), a cell septum/cytosolic protein (ZDS2, YALI0F12793g), an exportin transporter (CSE1, YALI0E07139g), and an uncharacterized protein (YALI0B18194g). One MG was downregulated in IL, a protein component of the polarisome (SPA2, YALI0F16665g), and one uncharacterized protein (YALI0F08107g) was classified with changed regulation in IL.

#### Enhanced IL-response by the mutant

Our gene classification of time series RNAseq data revealed 202 upregulated genes and 319 increasing genes for the MT strain in IL relative to the WT strain in IL (Figure 2D, H) (Supplementary File S3, data set iv). These genes were associated with lipid processes (i.e., membrane, membrane traffic protein, Golgi apparatus, endoplasmic reticulum), cell periphery (including cell cortex, cytoskeleton, cell wall organization/biogenesis and microtubule process), intracellular vesicles (i.e., vacuole, cytoplasmic vesicle, vesicle-mediated transport) and genetic processes (i.e., chromosome, chromatin/chromatin-binding/regulator and signal transduction) (Figure 2L). In contrast, the WT exhibited 327 upregulated genes and 223 increasing genes relative to the MT when both strains were cultured in IL. These genes were associated with mitochondrion, mitochondrial transmembrane transport, respirasome and the ribosome (Figure 2L). Interestingly, 8 of the MGs were identified in the classified genes. The MT upregulated an exportin transporter (CSE1, YALI0E07139g) and 3 genes were classified as increasing, including a chaperone (YALI0A09108g), a translation initiation factor (RPG1, YALI0C16247g), and protein component of the polarisome (SPA2, YALI0F16665g). Four of these MGs were classified with changed regulation including an actin cytoskeletal protein (YALI0A18381g), non-receptor serine/threonine protein kinase (CDC7, YALI0F23287g), and uncharacterized proteins (YALI0A13233g and YALI0C13002g).

### Identifying genotypes conferring IL tolerance

#### Choice of pair-wise data set to select gene targets conferring IL tolerance

In response to IL, there were 734 genes upregulated by the WT (Figure 2F) and 564 genes upregulated by the MT (Figure 2G). However, many of these differentially expressed genes are potentially a consequence of IL-toxicity (i.e., indirect effect) as opposed to conferring IL tolerance (i.e., direct effect). Therefore, using the appropriate pair-wise data set to select gene targets conferring IL tolerance is critical due to a large space of candidates and their combinations. Fortunately, the MT selected from ALE outperformed the WT when both were cultured in IL. It also possesses superior IL-tolerant genotypes. Taken altogether, we reasoned that, to identify genotypes conferring IL tolerance, the best gene candidates for reverse engineering high solvent tolerance could be found by comparing WT and MT gene expression in IL (Supplementary File S3, data set iv).

#### Formulation of Gene Co-Expression Connectivity (GeCCo) as a metric to select gene targets for reverse engineering

Conventionally, gene targets are selected based on greatest fold change between strains or conditions and/or gene ontology if the system is well understood. Using the pair-wise data set of WT and MT in IL, our analysis identified 202 candidate gene targets that were upregulated across 2 time points by the MT (Figure 2G). It is laborious to experimentally validate these genes for IL tolerance either in isolation or combinations. To reduce the most probable candidate list of gene targets for validation, we introduced GeCCo that selects highly connected and classified genes in the co-expression network.

In the first step of GeCCo, the co-expression network using the Pearson correlation (23) is built for the genes differentially expressed and regulated between the MT and WT strains in IL. Each gene in the co-expression network is classified using the gene classification methodology (see Materials and Methods, Figure S1). Figure 3A shows the network topology where each group of classified genes showed tight co-expression. For instance, genes that were upregulated or increased by the MT in IL were separated from genes that were upregulated or increased by the WT in IL. These gene clusters were connected by genes classified as changed regulation. In the second step, GeCCo selects the most connected genes in a subnetwork (e.g., upregulated gene subnetwork) based on their degree centrality. Here, degree centrality is the total number of edges connected to a vertex in the co-expression network. For instance, Figure 3B showed the most connected genes in the subnetwork of genes upregulated by the MT in IL.

**Figure 3.**
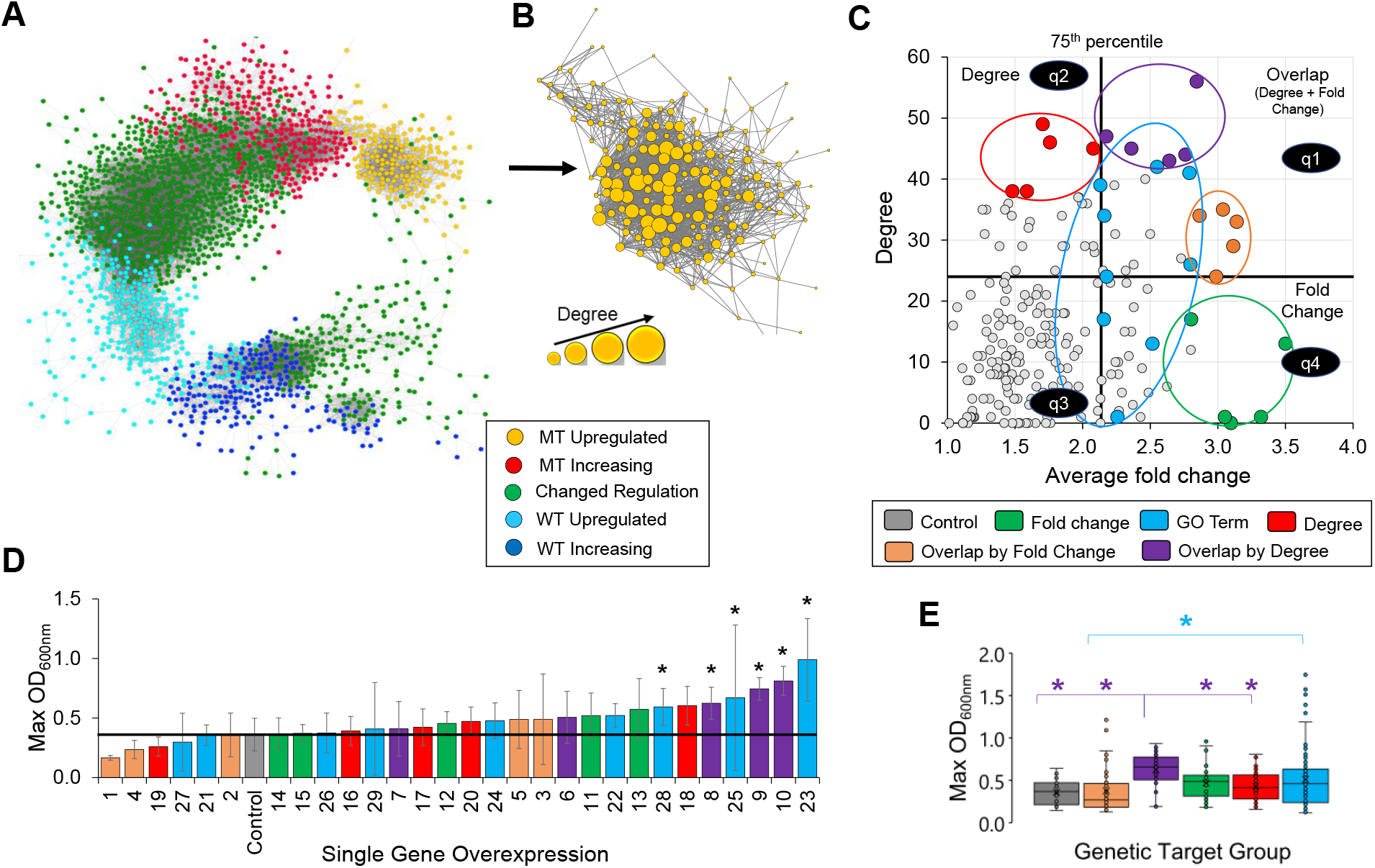
(**A**) Co-expression network of classified genes between the MT and WT in 8% IL. (**B**) Subnetwork of co-expressed genes upregulated by the MT in 8% IL illustrating gene connectivity by degree. (**C**) Gene targets selected by degree, fold change, overlap by fold change, overlap by degree, or gene ontology. (**D**) Maximum cell growth for individual overexpression of gene targets in 11% IL. (**E**) Box and whisker plot for each group of gene targets depicting all replicate maximum OD_600nm_ for all associated genes contained within each group. *p-value ≤ 0.05 (n ≥ 9 for each gene and n = 27 for the empty vector control).

#### Selection of gene targets by GeCCo for experimental validation

To demonstrate GeCCo, we used the subnetwork of genes upregulated by the MT in IL. For each of the 202 genes upregulated by the MT, the average fold change between the 2 transcriptomic time points (i.e., X (early) and Y (mid) scores, see Figure S1) and the degree of each gene from the co-expression subnetwork were calculated (Figure 3B). These genes were divided into 4 quadrants based on the 75^th^ percentile of average fold change and on the 75^th^ percentile of degree (Figure 3C). Quadrant 1 represents genes that were in the 75th percentile of both average fold change and degree, quadrant 2 represents genes that belong only in the 75^th^ percentile of degree, quadrant 3 represents genes that were not in either 75^th^ percentiles of degree or average fold change, and quadrant 4 represents genes that belong in both 75^th^ percentiles of degree and average fold change (Figure 3C). To test GeCCo, we selected the top 5 genes based on degree (quadrant 2), the top 5 genes based on degree overlapping with average fold change (i.e., overlap by degree, quadrant 1), and the top 5 genes based on average fold change overlapping with degree (i.e., overlap by fold change, quadrant 1). For control, we chose the top 5 genes based on average fold change (quadrant 4) and 9 genes based on gene ontology associations related to membrane, transport, kinase, cell wall, actin cytoskeleton organization, and myosin complex (quadrants 1, 2 and 4) (Figure 3C, Table S1).

### Validation of gene targets conferring IL tolerance

#### Single gene overexpression

Each of the 29 selected gene targets (Supplementary File S4) were tested for their efficacy in conferring IL tolerance by individually overexpressing each gene in the WT strain and characterizing cell growth in 11% IL (Figure 3D). In this extremely harsh concentration of IL, genes chosen from overlap by degree exhibited the best cell growth in IL among all the other target groups except genes chosen by GO term (Figure 3E). Of all the target groups, only 6 of the genes caused a decrease in cell mass including 3 genes chosen by overlap by fold change (#1, #2 and #4), 2 genes chosen by GO terms (#21 and #27), and 1 gene chosen by degree (#19).

Six genes exhibited a statistical increase in cell mass in this high concentration of IL (#23, #10, #9, #25, #8 and #28). Interestingly, these gene targets were selected from only 2 target groups of overlap by degree and GO association (Figure 3D). The 3 genes conferring enhanced IL tolerance chosen from overlap by degree included a membrane traffic protein (#8, TRS23, YALI0B22396g), a kinase activator (#9, CLB5, YALI0B15180g), and a microtubule binding motor protein (#10, KIP1/CIN8, YALI0F02673g). The 3 genes exhibiting increased IL-robustness selected by GO terms were annotated as an integral component of the membrane (#23, YALI0B12738g), a protein kinase (#25, SPS1, YALI0D19470g), and a myosin heavy chain (#28, MYO1, YALI0F13343g).

#### Dual-gene library enrichment

To identify the most IL-tolerant combination of genes, we duplicated the 29 gene target overexpression plasmids with a different selection marker (i.e., leucine and uracil selection markers) and combined the 2 sets of 29 overexpression plasmids into a library cocktail stock at equimolar concentrations (Figure 4A). This library was transformed into *Y. lipolytica* and 3 biological replicate transformants were cultured in 0%, 12% and 13% (v/v) IL for 2 rounds. While all strains grew quickly in 0% IL, cell growth was delayed for several days in the high IL concentrations (Figures 4B-4D). Interestingly, the dual-plasmid cocktail replicates grew faster than the empty-vector control in 12% IL (Figure 4C) and all replicates were able to grow in 13% IL while the empty-vector control showed no growth in this high concentration of IL (Figure 4D). Each cocktail replicate was sequenced after round-1 and round-2 cultures in 0%, 12% and 13% IL to identify enriched combinations of genes selected from the challenging pressure of IL (Supplementary File S5).

**Figure 4.**
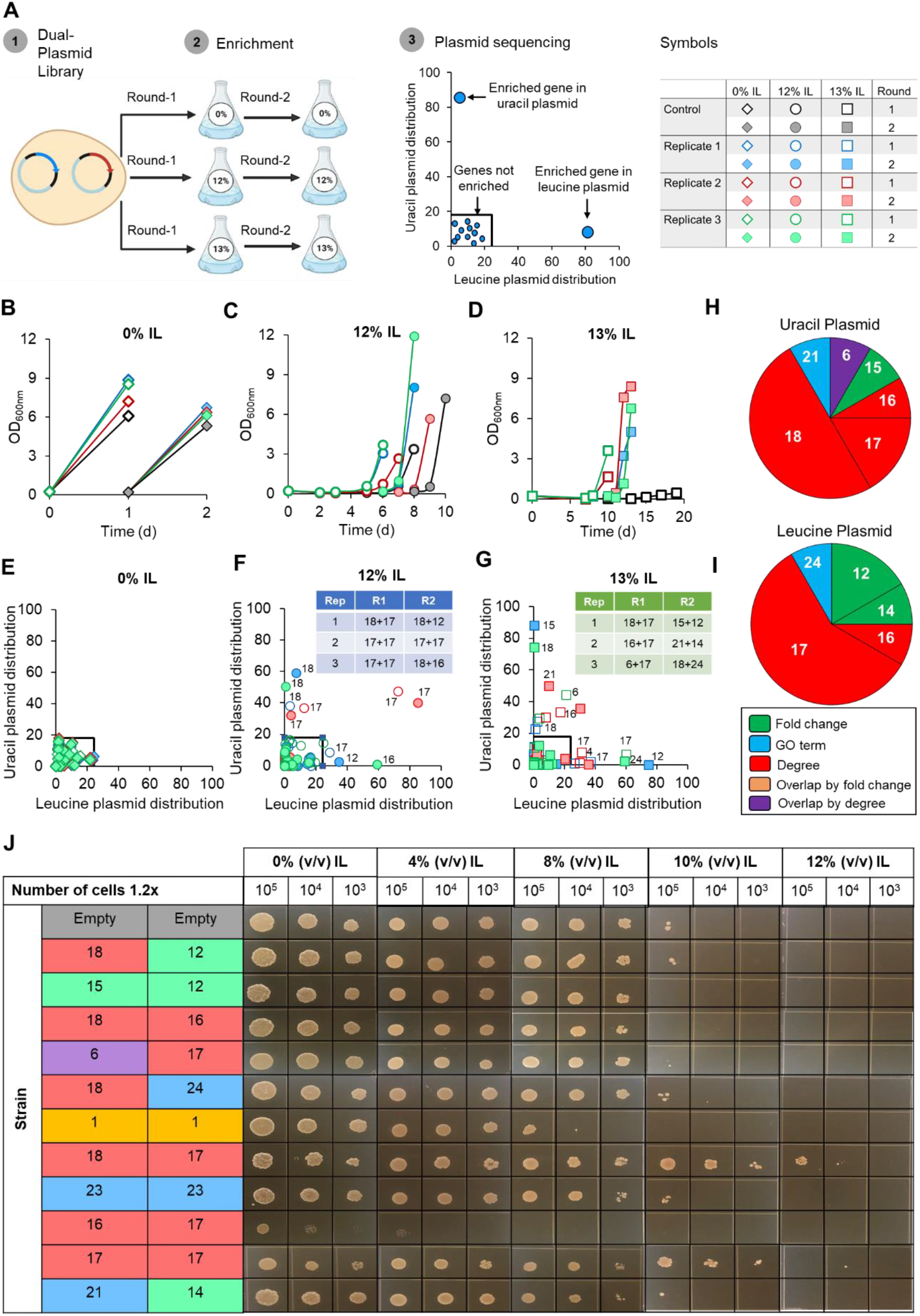
(**A**) Workflow of dual-gene library enrichment and plasmid sequencing (created with BioRender.com). Growth of empty-vector control and library cocktail replicates in (**B**) 0% (v/v) IL, (**C**) 12% (v/v) IL, and (**D**) 13% (v/v) IL. Plasmid sequencing to identify enriched combinations of genes for library-cocktail replicates in (**E**) 0% (v/v) IL, (**F**) 12% (v/v) IL, and (**G**) 13% (v/v) IL. (**H-I**) Frequency of enriched genes from library-cocktail enrichment selected by fold change (green), GO term (blue), degree (red), overlap by fold change (yellow), and overlap by degree (purple). (**J**) Plate spotting characterization of enriched combinations of genes from library-cocktail enrichment on 0%, 4%, 8%, 10%, and 12% (v/v) IL.

For all replicates cultured in 0% IL, genes were not enriched beyond 24.3% of the total gene pool in the leucine plasmid and 18% of the total gene pool in the uracil plasmid (Figure 4E). Plasmids sequenced from all replicate cultures in 12% and 13% IL revealed enriched combinations of genes (Figures 4F, G). Interestingly, replicate-2 cultured in 12% IL was the only replicate that maintained the same combination of genes between round-1 and round-2 (i.e., gene combination: #17+#17) (Figure 4F). However, gene combination #18+#17 was enriched in multiple replicates/rounds (i.e., 12% IL, replicate-1, round-1; 12% IL, replicate-3, round-1; and 13% IL, replicate-1, round-1). From these 9 combinations of genes, genes chosen by degree had the highest frequency in both the uracil plasmid (75% of genes) and leucine plasmid (66.7% of genes) (Figure 4H, 4I). Notably, genes chosen by fold change showed the second highest frequency (8.3% of genes in uracil plasmid; 25% of genes in leucine plasmid) followed by genes chosen by GO term (8.3% of genes in both uracil and leucine plasmids) (Figures 4H, 4I).

#### Characterization of enriched gene combinations for IL tolerance

Seeking to reverse engineer IL tolerance, we re-created strains individually bearing these 9 combinations of genes. For positive and negative controls, we generated strains bearing 2 copies of the best and worst genes from single overexpression characterization (i.e., #23+#23 and #1+#1) along with the empty-vector control. Remarkably, gene combination #18+#17 (that was enriched in multiple replicates) demonstrated significantly higher IL tolerance relative to all strains tested (Figure 4G, 4J). Interestingly, these genes were not directly co-expressed (i.e., not connected by an edge), but instead highly connected to other genes in the co-expression network. Gene ontology enrichment of gene #17 cluster (genes connected to #17) and gene #18 cluster (genes connected to #18), using GO-slim biological process, revealed both gene clusters were enriched in mitotic nuclear division (Figure 5C). Gene #17 cluster was also enriched for chromosome separation and mitotic chromosome condensation, microtubule cytoskeletal organization and microtubule-based movement, regulation of cyclin-dependent protein serine/threonine kinase activity, and protein phosphorylation (Figure 5C). For Gene #18 cluster, the only uniquely enriched term was Golgi vesicle transport (Figure 5C). The second-best strain contained the gene combination #17+#17 which was enriched in both rounds of 12% IL for replicate-2 (Figures 4F, 4J). Gene combination #1+#1, which showed increased sensitivity to IL in single-overexpression characterization, was the worst performing strain excluding gene combination #16+#17 which exhibited defective growth even when cultured without IL (Figure 4J). Taken together, dual-plasmid library enrichment revealed genes #18 and #17, both chosen by degree (i.e., connectivity), that conferred high IL tolerance when overexpressed simultaneously.

**Figure 5.**
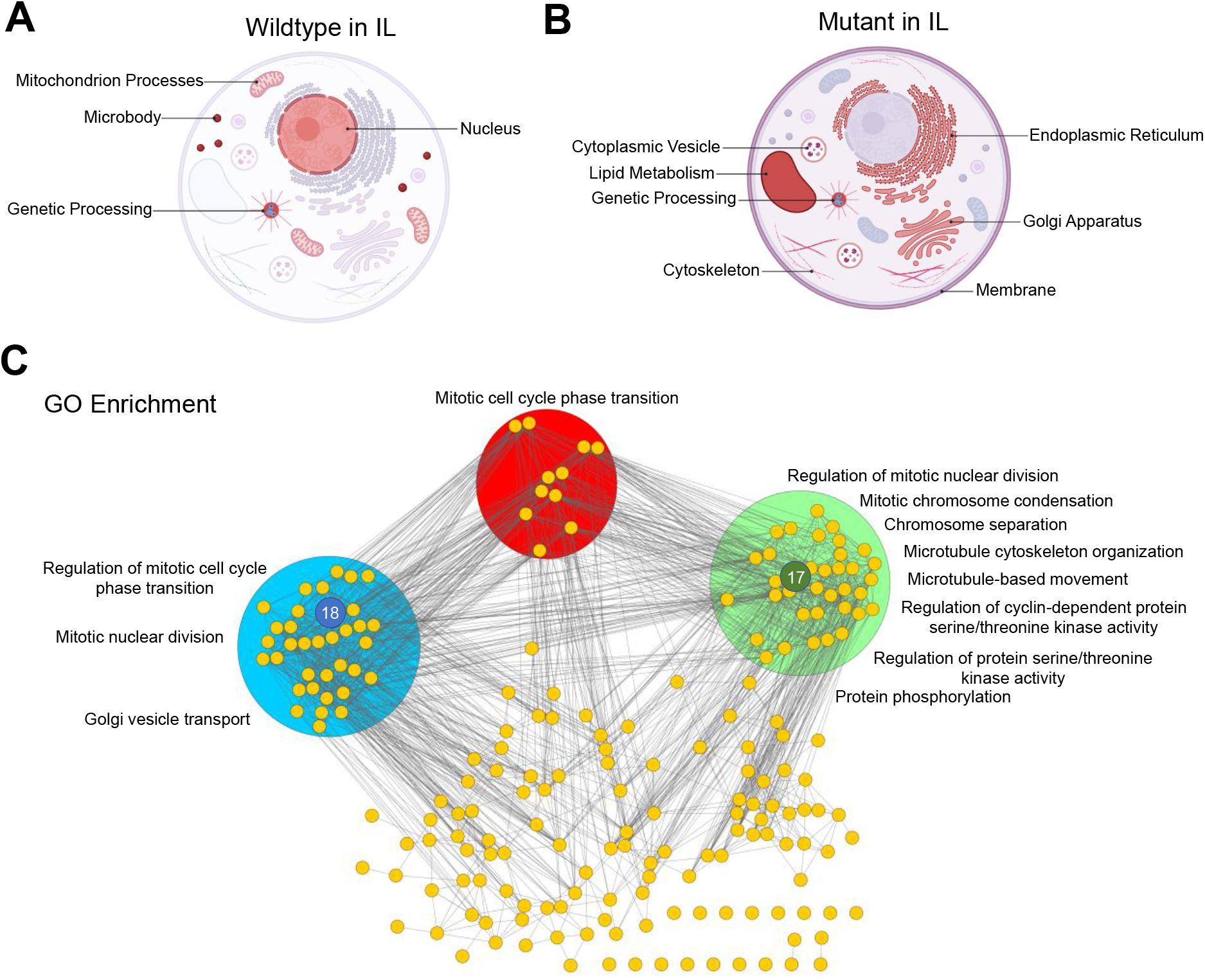
Cell model diagrams for (**A**) IL-response of WT, (**B**) IL-response of the MT, and (**C**) top performing genes when overexpressed individually (#8, #9, #10, #23, #25, and #28) and simultaneously (#17+#18) (created with BioRender.com).

## DISCUSSION

ILs inhibit microbial catalytic activity by affecting numerous cellular processes and components which makes IL tolerance a difficult phenotype to engineer. Currently, reverse engineering complex phenotypes requires extensive characterization to understand key mechanisms and often leads to numerous gene targets for further experimentation (24–26). Here, we developed GeCCo to identify a subset of promising gene targets for reverse engineering IL-robustness by exploiting a superior IL-tolerant MT strain’s gene co-expression connectivity. From the pool of genes upregulated by the MT in IL (relative to the WT in IL at both exponential time points), we discovered the best single gene targets (i.e., single gene overexpression) were predicted by gene connectivity overlapping in the 75^th^ percentile of fold change (i.e., overlap by degree) and the best combination of gene targets (i.e., dual gene overexpression) were predicted by co-expression connectivity alone (i.e., by degree).

Upon exposure to IL, only the WT increased expression of genes associated with mitochondrion processes (i.e., respirasome and mitochondrial transmembrane transport) (Figures 2J, 5A) suggesting ILs hinder mitochondrial function and oxidative phosphorylation. Interestingly, ILs interfered with ion transport across the mitochondrial membrane, negatively affecting mitochondria function and membrane potential in *S. cerevisiae* (27). In contrast, the WT repressed genes involved in cell wall organization/biogenesis (Figure 2J) which vindicates the reduced cell wall chitin content when *Y. lipolytica* is exposed to IL (22).

The MT strain, however, induced expression of genes associated with lipid metabolism in IL (e.g., membrane, endoplasmic reticulum, Golgi apparatus, membrane traffic proteins, cytoplasmic vesicle, etc.) (Figures 2K, 6B) which is supported by our previous discovery that biogenesis of sterols is critical for IL tolerance in *Y. lipolytica* (22). These results postulate other lipid metabolic processes (e.g., vesicle-mediated transport) underlying the superior IL tolerance of the MT strain as seen by that the drastic remodeling of lipid composition in IL (22). Both WT and MT strains responded to IL by increasing expression of genes associated with genetic processing (i.e., chromatin-binding/regulation, chromosome, transcription factors, and signal transduction) (Figures 2J, 2K, 5A, 5B) indicating ILs severely influence gene regulation and signal transduction.

Aiming to understand genotype(s) conferring high solvent tolerance, we reasoned that the best gene candidates would be genes upregulated by the MT relative to the WT when cultured in IL. Most of the gene targets, when individually overexpressed, increased cell mass in IL indicating our strategy of exploiting genes expressed more by the MT strain in this comparison was successful (Figure 3L), though we did not test genes from other pairwise comparisons (e.g., WT 8% IL vs. WT 0% IL). Strikingly, co-expression connectivity in conjunction with fold change (i.e., overlap by degree) had the highest statistical prediction of IL-tolerant genes when overexpressed individually (Figure 3E).

Even though the top 6 performing single gene targets have not previously been characterized in *Y. lipolytica*, these genes have orthologs in the model yeast *S. cerevisiae* except for gene #23. Gene #8 (TRS23, YALI0B22396g) is associated with regulation of traffic from endoplasmic reticulum to Golgi and autophagy (28). Gene #9 (CLB5, YALI0B15180g) is a non-essential B-type cyclin involved in replication of DNA (29) whereby deletion increases sensitivity to rapamycin, caffeine, hydroxyurea, doxorubicin, and many other chemicals (30–32). Gene #10 (CIN8, YALI0F02673g) is a non-essential kinesin motor protein with involvement in chromosome segregation and in mitotic spindle assembly (33). Notably, CIN8-deleted *S. cerevisiae* exhibited abnormally large cell size, poor growth, increased mitophagy and increased sensitivity to heat and various chemicals (34). Gene #25 (SPS1, YALI0D19470g) is a kinase regulator essential for prospore membrane closure (35). Gene #28 (MYO1, YALI0F13343g) is a class II myosin heavy chain with a key role in cytokinesis (36). Interestingly, deletion of MYO1 in S. *cerevisiae* decreases resistance to nikkomycin Z, and antibiotic that inhibits chitin synthase (37).

Interestingly, the dual-gene overexpression library enrichment revealed combinations of genes that differed from the top-performing individual overexpression results (Figures 4F, 4G). Most of the enriched gene targets in these dual-gene combinations were those selected by co-expression connectivity (i.e., degree) (Figures 4H, 4I), indicating inter-dependency between highly connected genes. This notion is further supported by the significant increase in IL tolerance when #17 and #18 were simultaneously overexpressed (Figure 4J) relative to the IL tolerance of either gene’s individual overexpression (Figure 3D).

Notably, these genes are not obvious targets to reverse engineer IL tolerance based on annotations alone. Gene #17 (BRN1, YALI0B03476g) is an essential gene in *S. cerevisiae* required for the mitotic chromosome segregation and condensation (38). Gene #18 (OYE2, YALI0D16247g) is a flavin mononucleotide oxidoreductase that mediates resistance to small alpha- and beta-unsaturated carbonyls such as acrolein, a product of lipid peroxidation (39) and increases abundance in response to DNA replication stress (40). Further, OYE2 (Gene #18) has been shown to be a powerful antioxidant protein in *S. cerevisiae* and overexpression of OYE2 (Gene #18) rescues Bax-induced cell death, a pro-apoptotic protein that causes hyperpolarization of the mitochondria membrane resulting in liberation of cytochrome c and reactive oxygen species (41). While lacking a strong connection as to how these two genes confer IL tolerance together, we can deduce that both genes are involved in the mutual process of mitotic cell cycle phase transition based on their co-expression gene clusters (Figure 5C). Further, we can reason that IL-toxicity is negatively affecting chromatin segregation during mitosis and proteins responsible for guiding and regulating these processes (i.e., microtubule and cytoskeletal organization, kinases, Golgi vesicle transport) are critical (Figure 5C).

While co-expression networks have been used to infer gene function, gene association and gene regulation, this is the first study where co-expression was used as a metric to select gene targets for reverse engineering to our knowledge. Though limited by the number of genes, phenotypes and organisms tested in our study, these results highlight GeCCo as a promising selection tool to predict gene targets for reverse engineering complex phenotypes.

## MATERIALS AND METHODS

### Strains

The *Y. lipolytica* strain, ATCC MYA-2613, a thiamine, leucine, and uracil auxotroph, was purchased from American Type Culture Collection and used as the parent strain (WT). The evolved strain YlCW001 (MT) was isolated after 200 generations in gradually increased concentrations of [EMIM][OAc] up to 18% (v/v) (22). The TOP10 *Escherichia coli* strain was used for molecular cloning. IL-gene overexpressing strains were generated by transforming constructed plasmids into WT via electroporation (42).

### Plasmids

The plasmid pSR001 (21), harboring leucine selection marker, TEF promoter and CYC1 terminator, was used as the backbone plasmid for individual IL-gene overexpression cassettes. The plasmid pSR008 (43), identical to pSR001 except the selection marker was replaced with uracil, was used for dual-gene overexpression. Both pSR001 and pSR008 were linearized by PCR amplification using primers 2 and 3 and each gene target was amplified from *Y. lipolytica* genomic DNA using the primers 79-136 listed in Table S2. Nucleotide sequences of the 29 target genes can be found in Supplementary File S4. Plasmids were constructed with Gibson assembly (44) using the linearized pSR001/pSR008 backbones and each amplified IL-gene. The complete list of strains and plasmids were documented in Table S2.

### Media and culturing conditions

#### Growth medium

Defined minimal media contained 6.7 g/L of yeast nitrogen base without amino acids (cat# Y0626, Sigma-Aldrich, MO, USA), 20 g/L of glucose, 380 mg/L leucine (cat# 172130250, Acros Organics, CA, USA), 76 mg/L uracil (cat# 157301000, Acros Organics, CA, USA), and various concentrations of [EMIM][OAc] (>95 % purity) (IoLiTec, AL, USA). Synthetic complete without leucine and uracil (SC-Leu-Ura) was prepared with 6.7 g/L of yeast nitrogen base without amino acids, 1.46 g/L of yeast synthetic drop-out medium supplement without uracil, leucine, and tryptophan (cat# Y1771, Sigma-Aldrich, MO, USA), 76 mg/L tryptophan (cat# 172110250, Acros Organics, CA, USA), 20 g/L glucose, and various concentrations of [EMIM][OAc]. SC without leucine (SC-Leu) was prepared by adding 76 mg/L uracil to SC-Leu-Ura media.

#### Culturing conditions

Seed cultures were conducted by inoculating a single colony from a fresh petri dish into 2 mL of media using a 14-mL culture tube and incubated overnight in a MaxQ6000 air incubator set at 28°C and 250 rpm. Sub-cultures were conducted by transferring 1 mL of seed cultures into 10 mL of media using 50 mL baffled flasks and cultured overnight until mid-exponential growth phase. Main experiment cultures were performed using 50 mL baffled flasks with 10 mL of media in 3 technical replicates per biological condition and repeated to achieve a minimum of 6 replicates.

#### Single gene overexpression characterization

Single-gene overexpression experiments were conducted using SC-Leu media containing 11% (v/v) [EMIM][OAc] with an inoculum of 0.25 OD_600nm_ from sub-seed cultures.

#### Dual-gene enrichment

Equimolar concentrations of all 29 gene overexpression plasmids with leucine selection marker and all 29 gene overexpression plasmids with uracil selection marker were combined into a library cocktail. The library cocktail was transformed into WT *Y. lipolytica* and recovered transformant colonies were combined and cultured in SC-Leu-Ura media. Round-1 of the enrichment experiment was conducted in 50 mL of SC-Leu-Ura media containing 0%, 12% or 13% (v/v) [EMIM][OAc] in 3 replicates alongside a strain bearing an empty-vector leucine plasmid and an empty-vector uracil plasmid (i.e., control). Once cells achieved an OD_600nm_ above 3, the culture broth was centrifuged, resuspended in water, and used to inoculate round-2 of enrichment before storing remaining cells at −80°C. Once cells achieved an OD_600nm_ above 3, round-2 cultures were immediately stored at −80°C. Plasmids were extracted after thawing cells to room temperature using the Invitrogen ChargeSwitch plasmid yeast mini kit (Fisher cat# CS10203).

#### Dual-gene plate screening

Dual-gene overexpression characterization was performed with SC-Leu-Ura media using sub-seed cultures that were centrifuged and resuspended to achieve an OD_600nm_ of 2 and diluted further by 10x and 100x. 2 µL of each dilution were transferred into gridded petri dishes containing SC-Leu-Ura media, 20 g/L agar and 0%, 4%, 8%, 10% or 12% (v/v) [EMIM][OAc] and incubated at 28°C for 2, 3, 5, 10 and 10 days for each [EMIM][OAc] concentration respectively. Plate screening was performed a minimum of 3 times for each construct.

### Analytical methods and bioinformatics

#### Variant detection (mutation analysis)

Whole genome resequencing reads of WT (GCA_009372015.1) and MT (GCA_009194645.1) strains were quality filtered as previously described (45) and imported to the CLC genomic workbench software version 11.0.1 (https://www.qiagenbioinformatics.com/). For each strain, the Map Reads to Refence tool was used with default parameters to map reads to reference genome Clib122 (GCA_000002525.1) (46). Variants for each strain were identified using the InDels and Structural Variants 1.81, Local Realignment 0.41, and Fixed Ploidy Variant Detection 1.67 tools all set with default parameters. Variants specific to the MT strain were identified using the Compare Sample Variant Tracks 1.4 tool and variants causing an amino acid change were identified using the Amino Acid Changes 2.4 tool. Types of nucleotide changes were classified in different categories including single nucleotide variances (SNV), multiple nucleotide variants (MNV), insertions, deletions, and replacements.

#### RNA-sequencing

Samples were collected in biological triplicates for RNA-sequencing from WT and MT strains at early- and mid-exponential growth phases in 0% and 8% (v/v) [EMIM][OAc]. Samples were immediately quenched in liquid nitrogen and stored at −80°C until RNA isolation. Total RNA was purified using the Qiagen RNeasy mini kit (cat #74104, Qiagen, CA, USA) prior to submission at the Joint Genome Institute (JGI) using Illumina sequencing. Filtered RNA-seq reads were analyzed within the CLC genomics workbench version 11.0.1 (https://www.qiagenbioinformatics.com/) using the RNA-seq analysis tool to produce transcripts per million (TPM) values for each gene/sample which was used for all downstream transcript-analyses (Supplementary File S6).

#### Plasmid amplification and Oxford Nanopore sequencing

Extracted plasmids were amplified using random hexamer primers (Thermo Scientific SO181) and φ29 DNA polymerase (NEB M0269L), as previously described (47), with the exception of using the commercial φ29 reaction buffer from NEB (B0269L). Amplification products were cleaned and concentrated using the E.Z.N.A.® Cycle-Pure Kit from Omega BioTek (D6492-02) and quantified using Nanodrop.

Purified DNA was then sequenced using Oxford Nanopore’s ligation sequencing chemistry (SQK-LSK109) in conjunction with native barcoding expansions (EXP-NBD104/114) for high throughput, according to the manufacturer’s recommendations. All samples were sequenced on a MinION device using R9.4.1 flowcells. Samples were demultiplexed and basecalled either i) in real time with MinKNOW v4.3.20 using the fast basecalling algorithm, or ii) asynchronously with Guppy GPU v5.0.11 on Ubuntu 20.04 LTS using the fast basecalling algorithm.

#### Plasmid sequence mapping

To call reads for certain gene/selection marker combinations in a sample, the demultiplexed FASTQ files were concatenated into a single file and run through Centrifuge (v 1.0.3) (48) in two iterations: i) using the uracil and leucine selection marker sequences as the “genomes” for alignment and then ii) using the 29 gene sequences as a basis for alignment (Supplementary File S4). Centrifuge then output .tsv files with each unique read ID that matched on the sequence provided, using the standard algorithm alignment cutoff parameters. The two output files were cross-referenced for reads that hit on both the selection marker and a gene. The number of reads for each combination of gene/selection marker was tabulated for each sample (Supplementary File S5). The number of reads for each gene was divided by the total number of reads (that mapped to genes and respective selection marker) for each sample’s uracil plasmid and leucine plasmid separately. This value was multiplied by 100 to represent the percentage of individual gene reads of the total reads for each plasmid in each sample.

#### Gene ontology enrichment

Gene ontology (GO) for genes containing variants identified from resequencing (i.e., mutated genes) were identified using ClueGo (49), integrated microbial genomes (IMG) (50) and blastP against the model yeast *S. cerevisiae* with E-value < 1 E-10. Annotations were combined and curated manually to classify mutated gene ontology. GO terms associated with classified genes were manually retrieved from the Panther database (51) (Supplementary File S7) and mapped to classified genes to calculate the number of genes associated with each annotation term. GO terms were defined as enriched if the sum of upregulated and increasing genes in an annotation term were equal to or greater than double the sum of upregulated and increasing genes between the two biological conditions in each pairwise data set. GO enrichment for gene clusters connected to gene #17 and gene #18 were conducted using the Panther overrepresentation test (http://geneontology.org/) (52–54) with the GO slim biological process database and default settings (Fisher’s Exact, Calculate false discovery rate), and parents terms were used for illustration.

#### Statistics

One-way analysis of variance (ANOVA) with Holm-Sidak correction was used for all statistical analyses with the SigmaPlot v.14 software.

### Gene Co-expression Connectivity (GeCCo) analysis

#### Gene expression classification

Gene TPM values were floored to a value of 5 and averaged into log_2_-scaled values to compare biological conditions (BC) in the following pairwise sets: i) MT 0% IL versus WT 0% IL (BC1), ii) WT 8% IL versus WT 0% IL (BC2), iii) MT 8% IL versus MT 0% IL (BC3), and iv) MT 8% IL versus WT 8% IL (BC4) (see Figure S1, Step 1). For each pairwise set, X score (i.e., early exponential fold change, equation 1), Y score (i.e., mid exponential fold change, equation 2) and Z score (i.e., genes that are regulated differently during exponential growth, equation 3) were calculated for each gene as shown below (also see Figure S1, Step 2):

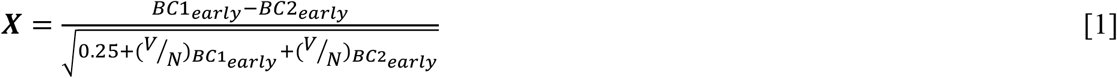

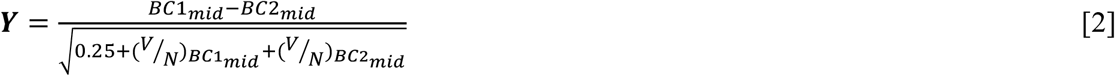

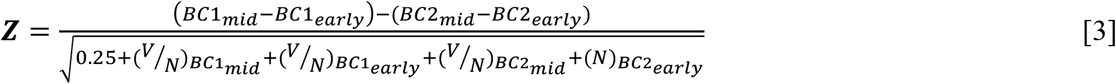

where BC1 and BC2 are average log_2_(TPM) values of a gene, subscripts early and mid are referred to early- and mid-exponential samples, V is variance, and N (=3) is the number of replicates. For each equation, the numerator calculates fold changes while the denominator accounts for error by dividing the fold change by the square root of pseudo variance (0.25) and the summation of variance (V) divided by respective number of replicates for each sample in the numerator.

Next, genes were categorized into 1 of 6 classes based on the values of X, Y and Z scores, including : BC1 upregulated (X ≥ 1 and Y ≥ 1), BC1 increasing (−1 < X < 1, Y ≥ 1, and Z ≥ 1.5), BC2 upregulated (X ≤ −1, Y ≤ −1), BC2 increasing (−1 < X < 1, Y ≤ −1, and Z ≤ −1.5), changed regulation (|Z| ≥ 1.5), and no change (Figure S1, Step 3). The “upregulated” gene class represents genes with greater expression values at both early- and mid-exponential samples. The “increasing” gene class represents genes without a significant change in expression value at the early-exponential sample but a greater expression value at mid-exponential sample. The “changed regulation” gene class represents genes that are neither upregulated nor increasing but with a regulation score greater than 1.5. The “no change” gene class represents genes that fail to meet the requirements of upregulated, increasing or changed regulation classifications.

#### Gene co-expression network construction and gene target selection

Pearson correlation was conducted on early- and mid-exponential WT 8% IL and MT 8% IL sample-gene TPM values using a stringent cutoff of 0.95. Next, this co-expression network was reduced by removing all genes except those classified as upregulated by the MT 8% IL in pairwise set iv (i.e., MT 8%IL vs WT 8%IL) and used to calculate degree centrality. These genes were ranked according to their degree and their average fold change between the two transcriptomic time points (i.e., average of X and Y scores) and the 75^th^ percentiles were calculated for both criteria. Finally, we chose 29 gene targets from the top 5 genes ranked by fold change, the top 5 genes by degree, the top 5 genes by degree overlapping with fold change, the top 5 genes by fold change overlapping with degree, and 9 genes with relevant gene ontology terms (e.g., membrane, kinase, transport, cell wall and myosin complex). GeCCo can be found at https://github.com/TrinhLab/GeCCo/.

## ACKNOWLEDGEMENTS

The authors would like to acknowledge financial support from the DOE BER Genomic Science Program award DE-SC0019412. The genome resequencing and RNAseq work was supported by the DOE FICUS #50384 award, which was performed at the U.S. DOE Joint Genome Institute (JGI). JGI is a DOE Office of Science User Facility, that is supported by the Office of Science of the U.S. DOE under Contract No. DE-AC02-05CH11231. The views, opinions, and/or findings contained in this article are those of the authors and should not be interpreted as representing the official views or policies, either expressed or implied, of the funding agencies.

## AUTHOR CONTRIBUTIONS

CT conceived and supervised the study. CW, SR, SG, DD, BM, and CT designed the experiments, and analyzed the data. CW, SR, SG, DD, and BM performed the experiments. CW and CT wrote the manuscript with the co-authors’ inputs. All authors read and approved the manuscript.

## SUPPLEMENTARY DATA

**Supplementary File S1** contains Tables S1, S2, S3 and Figure S1. **Table S1:** Gene target locus tag, accession, average fold change, degree, category of selection, and ortholog of *Saccharomyces cerevisiae*. **Table S2:** Primers used for constructing gene overexpression plasmids. **Table S3:** Strains and plasmids used for single- and dual-gene overexpression. **Supplementary Figure S1:** Gene classification methodology.

**Supplementary File S2:** A list of genetic mutations of the evolved strain YlCW001.

**Supplementary File S3:** Gene classification for i) mutant in 0% IL versus wildtype in 0% IL, ii) wildtype in 0% IL versus wildtype in 8%, iii) mutant in 0% IL versus mutant in 8% IL, and iv) mutant in 8% IL versus wildtype in 8% IL.

**Supplementary File S4:** Nucleotide sequences of target genes.

**Supplementary File S5:** Target dual-gene enrichment.

**Supplementary File S6:** TPM values for both WT and MT *Y. lipolytica* strains growing in 0% and 8% IL.

**Supplementary File S7:** Panther GO terms.

## REFERENCES

1. P.-Y. Kim, D. J. Pollard, J. M. Woodley, Substrate Supply for Effective Biocatalysis. Biotechnology Progress 23, 74–82 (2007).

2. A. M. Klibanov, Improving enzymes by using them in organic solvents. Nature 409, 241–246 (2001).

3. L. J. Bruce, A. J. Daugulis, Solvent Selection Strategies for Extractive Biocatalysis. Biotechnology Progress 7, 116–124 (1991).

4. T. Itoh, “Biotransformation in ionic liquid” in Future directions in biocatalysis. (Elsevier, 2017), pp. 27–67.

5. M. Moniruzzaman, K. Nakashima, N. Kamiya, M. Goto, Recent advances of enzymatic reactions in ionic liquids. Biochemical Engineering Journal 48, 295–314 (2010).

6. C. Li et al., Comparison of dilute acid and ionic liquid pretreatment of switchgrass: Biomass recalcitrance, delignification and enzymatic saccharification. Bioresource Technology 101, 4900–4906 (2010).

7. C. G. Yoo, Y. Pu, A. J. Ragauskas, Ionic liquids: Promising green solvents for lignocellulosic biomass utilization. Current Opinion in Green and Sustainable Chemistry 5, 5–11 (2017).

8. A. M. Socha et al., Efficient biomass pretreatment using ionic liquids derived from lignin and hemicellulose. Proceedings of the National Academy of Sciences 111, E3587 (2014).

9. E. T. Mohamed et al., Generation of a platform strain for ionic liquid tolerance using adaptive laboratory evolution. Microbial Cell Factories 16, 204 (2017).

10. M. Frederix et al., Development of an E. coli strain for one-pot biofuel production from ionic liquid pretreated cellulose and switchgrass. Green Chemistry 18, 4189–4197 (2016).

11. Y.-W. Wu et al., Ionic Liquids Impact the Bioenergy Feedstock-Degrading Microbiome and Transcription of Enzymes Relevant to Polysaccharide Hydrolysis. mSystems 1 (2016).

12. M. Ouellet et al., Impact of ionic liquid pretreated plant biomass on Saccharomyces cerevisiae growth and biofuel production. Green Chemistry 13, 2743–2749 (2011).

13. M. Yu et al., Effects of the 1-alkyl-3-methylimidazolium bromide ionic liquids on the antioxidant defense system of Daphnia magna. Ecotoxicology and environmental safety 72, 1798–1804 (2009).

14. K. M. Docherty, J. C. F. Kulpa, Toxicity and antimicrobial activity of imidazolium and pyridinium ionic liquids. Green Chemistry 7, 185–189 (2005).

15. L.-P. Liu et al., Mechanistic insights into the effect of imidazolium ionic liquid on lipid production by Geotrichum fermentans. Biotechnology for Biofuels 9, 266 (2016).

16. H. G. Lim et al., Generation of ionic liquid tolerant Pseudomonas putida KT2440 strains via adaptive laboratory evolution. Green Chemistry 22, 5677–5690 (2020).

17. T. L. Ruegg et al., An auto-inducible mechanism for ionic liquid resistance in microbial biofuel production. Nature communications 5, 3490 (2014).

18. D. A. Higgins et al., Natural Variation in the Multidrug Efflux Pump SGE1 Underlies Ionic Liquid Tolerance in Yeast. Genetics 210, 219–234 (2018).

19. K. B. Reed, J. M. Wagner, S. d’Oelsnitz, J. M. Wiggers, H. S. Alper, Improving ionic liquid tolerance in Saccharomyces cerevisiae through heterologous expression and directed evolution of an ILT1 homolog from Yarrowia lipolytica. Journal of Industrial Microbiology and Biotechnology 46, 1715–1724 (2019).

20. I. R. Sitepu et al., Yeast tolerance to the ionic liquid 1-ethyl-3-methylimidazolium acetate. FEMS Yeast Research 14, 1286–1294 (2014).

21. S. Ryu, N. Labbé, C. T. Trinh, Simultaneous saccharification and fermentation of cellulose in ionic liquid for efficient production of α-ketoglutaric acid by Yarrowia lipolytica. Applied Microbiology and Biotechnology 99, 4237–4244 (2015).

22. C. Walker, S. Ryu, C. T. Trinh, Exceptional solvent tolerance in Yarrowia lipolytica is enhanced by sterols. Metabolic Engineering 54, 83–95 (2019).

23. V. Randhawa, S. Pathania, Advancing from protein interactomes and gene co-expression networks towards multi-omics-based composite networks: approaches for predicting and extracting biological knowledge. Briefings in Functional Genomics 19, 364–376 (2020).

24. Y. Chen et al., Reverse engineering of fatty acid-tolerant Escherichia coli identifies design strategies for robust microbial cell factories. Metabolic Engineering 61, 120–130 (2020).

25. R. Mans, J.-M. G. Daran, J. T. Pronk, Under pressure: evolutionary engineering of yeast strains for improved performance in fuels and chemicals production. Current Opinion in Biotechnology 50, 47–56 (2018).

26. K. R. Choi et al., Systems Metabolic Engineering Strategies: Integrating Systems and Synthetic Biology with Metabolic Engineering. Trends in Biotechnology 37, 817–837 (2019).

27. Q. Dickinson et al., Mechanism of imidazolium ionic liquidstoxicity in Saccharomyces cerevisiae and rationalengineering of a tolerant, xylose-fermenting strain. Microbial Cell Factories 15, 17 (2016).

28. M. Sacher, J. Barrowman, D. Schieltz, J. R. Yates, S. Ferro-Novick, Identification and characterization of five new subunits of TRAPP. European Journal of Cell Biology 79, 71–80 (2000).

29. E. Schwob, K. Nasmyth, CLB5 and CLB6, a new pair of B cyclins involved in DNA replication in Saccharomyces cerevisiae. Genes & Development 7, 1160–1175 (1993).

30. A. M. Dudley, D. M. Janse, A. Tanay, R. Shamir, G. M. Church, A global view of pleiotropy and phenotypically derived gene function in yeast. Mol Syst Biol 1, 2005.0001-2005.0001 (2005).

31. T. J. Westmoreland et al., Comparative Genome-Wide Screening Identifies a Conserved Doxorubicin Repair Network That Is Diploid Specific in Saccharomyces cerevisiae. PLOS ONE 4, e5830 (2009).

32. L. Kapitzky et al., Cross-species chemogenomic profiling reveals evolutionarily conserved drug mode of action. Mol Syst Biol 6, 451 (2010).

33. M. A. Hoyt, L. He, K. K. Loo, W. S. Saunders, Two Saccharomyces cerevisiae kinesin-related gene products required for mitotic spindle assembly. J Cell Biol 118, 109–120 (1992).

34. F. Basmaji et al., The ‘interactome’ of the Knr4/Smi1, a protein implicated in coordinating cell wall synthesis with bud emergence in Saccharomyces cerevisiae. Molecular Genetics and Genomics 275, 217–230 (2006).

35. M. A. Iwamoto, S. R. Fairclough, S. A. Rudge, J. Engebrecht, Saccharomyces cerevisiae Sps1p regulates trafficking of enzymes required for spore wall synthesis. Eukaryot Cell 4, 536–544 (2005).

36. F. Z. Watts, G. Shiels, E. Orr, The yeast MYO1 gene encoding a myosin-like protein required for cell division. EMBO J 6, 3499–3505 (1987).

37. F. E. Rivera-Molina, S. González-Crespo, Y. Maldonado-De la Cruz, J. M. Ortiz-Betancourt, J. R. Rodríguez-Medina, 2,3-Butanedione monoxime increases sensitivity to Nikkomycin Z in the budding yeast Saccharomyces cerevisiae. World J Microbiol Biotechnol 22, 255–260 (2006).

38. B. D. Lavoie, K. M. Tuffo, S. Oh, D. Koshland, C. Holm, Mitotic chromosome condensation requires Brn1p, the yeast homologue of Barren. Mol Biol Cell 11, 1293–1304 (2000).

39. W. Trotter Eleanor, J. Collinson Emma, W. Dawes Ian, M. Grant Chris, Old Yellow Enzymes Protect against Acrolein Toxicity in the Yeast Saccharomyces cerevisiae. Applied and Environmental Microbiology 72, 4885–4892 (2006).

40. J. M. Tkach et al., Dissecting DNA damage response pathways by analysing protein localization and abundance changes during DNA replication stress. Nature Cell Biology 14, 966–976 (2012).

41. O. Odat et al., Old Yellow Enzymes, Highly Homologous FMN Oxidoreductases with Modulating Roles in Oxidative Stress and Programmed Cell Death in Yeast*. Journal of Biological Chemistry 282, 36010–36023 (2007).

42. K. A. Markham, S. Vazquez, H. S. Alper, High-efficiency transformation of Yarrowia lipolytica using electroporation. FEMS Yeast Research 18 (2018).

43. S. Ryu, T. Trinh Cong, A. Elliot Marie, Understanding Functional Roles of Native Pentose-Specific Transporters for Activating Dormant Pentose Metabolism in Yarrowia lipolytica. Applied and Environmental Microbiology 84, e02146–02117.

44. D. G. Gibson et al., Enzymatic assembly of DNA molecules up to several hundred kilobases. Nature Methods 6, 343–345 (2009).

45. C. Walker et al., Draft Genome Assemblies of Ionic Liquid-Resistant Yarrowia lipolytica PO1f and Its Superior Evolved Strain, YlCW001. Microbiology Resource Announcements 9, e01356–01319 (2020).

46. B. Dujon et al., Genome evolution in yeasts. Nature 430, 35–44 (2004).

47. F. B. Dean et al., Comprehensive human genome amplification using multiple displacement amplification. Proceedings of the National Academy of Sciences 99, 5261 (2002).

48. D. Kim, L. Song, F. P. Breitwieser, S. L. Salzberg, Centrifuge: rapid and sensitive classification of metagenomic sequences. Genome Res 26, 1721–1729 (2016).

49. G. Bindea et al., ClueGO: a Cytoscape plug-in to decipher functionally grouped gene ontology and pathway annotation networks. Bioinformatics 25, 1091–1093 (2009).

50. I. M. A. Chen et al., The IMG/M data management and analysis system v.6.0: new tools and advanced capabilities. Nucleic Acids Research 49, D751–D763 (2021).

51. H. Mi et al., PANTHER version 16: a revised family classification, tree-based classification tool, enhancer regions and extensive API. Nucleic Acids Research 49, D394–D403 (2021).

52. M. Ashburner et al., Gene Ontology: tool for the unification of biology. Nature Genetics 25, 25–29 (2000).

53. C. Gene Ontology, The Gene Ontology resource: enriching a GOld mine. Nucleic acids research 49, D325–D334 (2021).

54. H. Mi, A. Muruganujan, D. Ebert, X. Huang, P. D. Thomas, PANTHER version 14: more genomes, a new PANTHER GO-slim and improvements in enrichment analysis tools. Nucleic acids research 47, D419–D426 (2019).

