## Supplementary File S1 for "Gene Co-expression Connectivity Predicts Gene Targets Underlying High Ionic Liquid Tolerance in *Yarrowia lipolytica*"

### Gene Co-expression Connectivity Predicts Gene Targets Underlying High Ionic Liquid Tolerance in *Yarrowia lipolytica*

Caleb Walker, Seunghyun Ryu, Sergio Garcia, David Dooley, Brian Mendoza, Cong T. Trinh*

Department of Chemical and Biomolecular Engineering, University of Tennessee, Knoxville, TN 37996

**Supplementary Table S1.** Gene target locus tag, accession, average fold change, degree, category of selection, and ortholog of *Saccharomyces cerevisiae*.

| Gene | Locus tag | Accession | Average fold change | Degree | Category | *S. cerevisiae* Orthologue |
|---|---|---|---|---|---|---|
| 1 | YALI0E10417g | Q6C6D3 | 3.14 | 33 | Overlap by fold change | N/A |
| 2 | YALI0C21758g | Q6CB57 | 3.11 | 29 | Overlap by fold change | KIN4 |
| 3 | YALI0B16324g | Q6CEE3 | 3.04 | 35 | Overlap by fold change | CHS2 |
| 4 | YALI0D08118g | Q6C9V3 | 2.99 | 24 | Overlap by fold change | N/A |
| 5 | YALI0E00792g | Q6C7H5 | 2.86 | 34 | Overlap by fold change | KAR9 |
| 6 | YALI0F06402g | Q6C2N4 | 2.84 | 56 | Overlap by degree | YCS4 |
| 7 | YALI0E07491g | Q6C6P8 | 2.17 | 47 | Overlap by degree | KIP3 |
| 8 | YALI0B22396g | Q6CDP3 | 2.36 | 45 | Overlap by degree | TRS23 |
| 9 | YALI0B15180g | Q6CEJ4 | 2.76 | 44 | Overlap by degree | CLB5 |
| 10 | YALI0F02673g | Q6C348 | 2.64 | 43 | Overlap by degree | KIP1/CIN8 |
| 11 | YALI0C03377g | Q6CD67 | 3.50 | 13 | Fold Change (FC) | CDC20 |
| 12 | YALI0E12419g | Q6C650 | 3.32 | 1 | Fold Change (FC) | N/A |
| 13 | YALI0E21549g | Q6C529 | 3.10 | 0 | Fold Change (FC) | BIO3 |
| 14 | YALI0E17545g | Q6C5J6 | 3.05 | 1 | Fold Change (FC) | N/A |
| 15 | YALI0C21296g | Q6CB77 | 2.80 | 17 | Fold Change (FC) | N/A |
| 16 | YALI0A18722g | F2Z6B2 | 1.70 | 49 | Degree | VPS28 |
| 17 | YALI0B03476g | Q6CFV1 | 1.76 | 46 | Degree | BRN1 |
| 18 | YALI0D16247g | Q6C8X0 | 2.08 | 45 | Degree | OYE2 |
| 19 | YALI0E34852g | Q6C3H2 | 1.48 | 38 | Degree | EMP24 |
| 20 | YALI0F27423g | Q6C066 | 1.59 | 38 | Degree | N/A |
| 21 | YALI0D12232g | Q6C9C8 | 2.79 | 41 | GO Term | N/A |
| 22 | YALI0C08096g | Q6CCM9 | 2.80 | 26 | GO Term | N/A |
| 23 | YALI0B12738g | Q6CEV0 | 2.16 | 34 | GO Term | N/A |
| 24 | YALI0C18183g | Q6CBJ7 | 2.16 | 17 | GO Term | N/A |
| 25 | YALI0D19470g | Q6C8H9 | 2.55 | 42 | GO Term | SPS1 |
| 26 | YALI0C21802g | Q6CB55 | 2.52 | 13 | GO Term | SEC27 |
| 27 | YALI0B09273g | Q6CF88 | 2.26 | 1 | GO Term | VCX1 |
| 28 | YALI0F13343g | Q6C1U3 | 2.18 | 24 | GO Term | MYO1 |
| 29 | YALI0E29557g | Q6C455 | 2.13 | 39 | GO Term | HOF1 |

**Supplementary Table S2.** Primers used for constructing IL gene overexpression plasmids.

| Gene ID | Locus Tag | Primer | Sequence |
|---|---|---|---|
| pSR001 | N/A | 2 | TTTGAATGATTCTTATACTCAGAAG |
| | N/A | 3 | TCATGTAATTAGTTATGTCACGCTTAC |
| #1 | YALI0E10417g | 79 | ccttctgagtataagaatcattcaaaATGCGCGCCATTGAGGGCCTGGGC |
| | | 91 | gcgtgacataactaattacatgaTTAGAAACCGGTGAAGCCCGACTGCATGGAG |
| #2 | YALI0C21758g | 80 | ccttctgagtataagaatcattcaaaATGTCTCGACAAAAAAAATCGCGAGAAATCCGG |
| | | 92 | gcgtgacataactaattacatgaTTAAGATGCAGAAACTCTTGACCGTCGTCGG |
| #3 | YALI0B16324g | 81 | ccttctgagtataagaatcattcaaaATGTCGAACAAGCCTCGAGACCTGCCCGAGCC |
| | | 93 | gcgtgacataactaattacatgaTTACTGTTGGCCAGTGAGCCCCTGGTGTCTTTTGAGG |
| #4 | YALI0D08118g | 82 | ccttctgagtataagaatcattcaaaATGCCAATATTTGGGTATTACCGCTTTGGG |
| | | 94 | gcgtgacataactaattacatgaTTACAATCTCTCAACAGGCCCCTCGAAATCAGCCTC |
| #5 | YALI0E00792g | 83 | ccttctgagtataagaatcattcaaaATGCTATCCACGTGTGCTTCCAATCTCGAGAAGC |
| | | 95 | gcgtgacataactaattacatgaCTACCGGACCACTGGTTTGGCGATACGCACG |
| #6 | YALI0F06402g | 84 | ccttctgagtataagaatcattcaaaATGTCGGAATTCATTCTCAACGAGGTGTACACAG |
| | | 96 | gcgtgacataactaattacatgaTTAGACCTTACCCACCACCTTGAAACCCTCGGC |
| #7 | YALI0E07491g | 85 | ccttctgagtataagaatcattcaaaATGGAGTCTTCTATTTCAGTGGCCGTCAGAGTAC |
| | | 97 | gcgtgacataactaattacatgaCTACTGTGTCACCCTCTGGGGCATCCCCAGC |
| #8 | YALI0B22396g | 86 | ccttctgagtataagaatcattcaaaATGACCGTGTATGCACTCTACATTCTCAACAAAG |
| | | 98 | gcgtgacataactaattacatgaCTAGCTCACCGACTCGAGGTACTTGTTGAGATGCAGG |
| #9 | YALI0B15180g | 87 | ccttctgagtataagaatcattcaaaATGAACTTTCCAGTGAGTACTAATGACCCGGAC |
| | | 99 | gcgtgacataactaattacatgaTTAGATAAATAGGTTAGTTTCGTAAGAATTGACACC |
| #10 | YALI0F02673g | 88 | ccttctgagtataagaatcattcaaaATGTCTTCGCGCCACAACAGTTTGGGAATGAGG |
| | | 100 | gcgtgacataactaattacatgaCTACTGCTTTCTATACTTCCTCGGAGGCACCAACCTG |
| #11 | YALI0C03377g | 89 | ccttctgagtataagaatcattcaaaATGTCGACAAAATCGCCTGTCACTCCTCGATACG |
| | | 101 | gcgtgacataactaattacatgaCTATCTAATACCCACAGTGGAAATCTCCTTGCTCTGG |
| #12 | YALI0E12419g | 90 | ccttctgagtataagaatcattcaaaATGACACAGATTATCCATAACGCCACCATCCCC |
| | | 102 | gcgtgacataactaattacatgaTTACAGCTTGGCCTTGTTGAGACCCAGAACAACATCG |

| | | | |
|---|---|---|---|
| #13 | YALI0E21549g | 103 | ccttctgagtataagaatcattcaaaATGAAAATCGGCGGTCTGTGGAAAAAATTCCCCG |
| | | 115 | gcgtgacataactaattacatgaTTACTGAAAGTAAGTTCTCACTCTCTGTTGAATGGC |
| #14 | YALI0E17545g | 104 | ccttctgagtataagaatcattcaaaATGCTCAAGCAACTGGAAAATATTCTTGAGTCGC |
| | | 116 | gcgtgacataactaattacatgaCTACAACCGCCCTCTCATCTCCTTCTGAGAATGCC |
| #15 | YALI0C21296g | 105 | ccttctgagtataagaatcattcaaaATGGACACTCCGACCCGTCCCCGAGCCGATCTG |
| | | 117 | gcgtgacataactaattacatgaCTACACCTTCAGTTTGGCCTCAAGCTTCATAAAATCC |
| #16 | YALI0A18722g | 106 | ccttctgagtataagaatcattcaaaATGTCACTTCCATACGCCCCACTAACCCATATGC |
| | | 118 | gcgtgacataactaattacatgaCTAATCTAGTTCTGCAAAGAAACCCTTGTAAG |
| #17 | YALI0B03476g | 107 | ccttctgagtataagaatcattcaaaATGGCAAGACCAGCTCGACGACGGTCTTCCGG |
| | | 119 | gcgtgacataactaattacatgaTTATGCTCCCAGCTGATCCAGGCGAACATTCATATCC |
| #18 | YALI0D16247g | 108 | ccttctgagtataagaatcattcaaaATGACACAAACGCACAATCTGTTTTCGCCAATC |
| | | 120 | gcgtgacataactaattacatgaTTACATCTTGTACGCAGGGTAATCAGTGTAGCCC |
| #19 | YALI0E34852g | 109 | ccttctgagtataagaatcattcaaaATGAAGTTCCTCGTTTGTCTTCTGGTTCTGGCGC |
| | | 121 | gcgtgacataactaattacatgaTTAAACCAAAGTCTTGACCTCAAAGAATCGC |
| #20 | YALI0F27423g | 110 | ccttctgagtataagaatcattcaaaATGTTTGGCATCTTTTCGCGCATTCCCGTGGAGC |
| | | 122 | gcgtgacataactaattacatgaCTAACTGCTCGTAGTGGCCACCTCAACAGGCTCG |
| #21 | YALI0D12232g | 111 | ccttctgagtataagaatcattcaaaATGATCATCAGCCTGACTGTGACCGTTAAAACG |
| | | 123 | gcgtgacataactaattacatgaTCAAACAACCTCTTCTCGGACAAAGCCCCGGC |
| #22 | YALI0C08096g | 112 | ccttctgagtataagaatcattcaaaATGGAGCACTTCACTGCGCACGATGGGTCGTTCC |
| | | 124 | gcgtgacataactaattacatgaTTACATAATAACACAATCGTCGGCCTCCTTGGTCACC |
| #23 | YALI0B12738g | 113 | ccttctgagtataagaatcattcaaaATGGACACGAAACGAAGCCTTTCAGGGGCTCCCC |
| | | 125 | gcgtgacataactaattacatgaCTACCTCGTCAGACCAACTCCTAATCCTACCG |
| #24 | YALI0C18183g | 114 | ccttctgagtataagaatcattcaaaATGTTGCCACGAGCCCTGAGTCGCCACGTGG |
| | | 126 | gcgtgacataactaattacatgaCTACTTATAGTCTGCGACTTCTTCAACGTCACCAGGG |
| #25 | YALI0D19470g | 127 | ccttctgagtataagaatcattcaaaATGGCATCGGCAAGCTCAGAACTATCACGGCTGG |

| | | | |
|---|---|---|---|
| | | 132 | gacataactaattacatgaCTAGTAACGATCGTACTCTTCGAACTCCTCGATCCACCG |
| #26 | YALI0C21802g | 128 | ccttctgagtataagaatcattcaaaATGAAGCTCGAAATCAAGGTGAGTAATTGAGCG |
| | | 133 | gcgtgacataactaattacatgaTTACAAGGTGTTGGCAGCCTCATCAACCTCCTCC |
| #27 | YALI0B09273g | 129 | ccttctgagtataagaatcattcaaaATGCCTTCCGAATCCACCCCTCTCATTGGCCGAC |
| | | 134 | gcgtgacataactaattacatgaTTACATATCCGAGGGATAGTAGAAGAAGGCAATGGCG |
| #28 | YALI0F13343g | 130 | ccttctgagtataagaatcattcaaaATGTGCGTGATGATGTTTTGTTTCAGCTGGATTG |
| | | 135 | gcgtgacataactaattacatgaTTAAACAAAGGAGGGCTCAAAGTTGGAGTTGTTGTTG |
| #29 | YALI0E29557g | 131 | ccttctgagtataagaatcattcaaaATGTCGGAACATTCGTTTGCCAACAACTTTTGGG |
| | | 136 | gcgtgacataactaattacatgaCTAAATGTCAGCCAAAAAGTTGCTGGGAGCTAGACC |

**Supplementary Table S3:** Strains and plasmids used for single- and dual-gene overexpression experiments. Strain #i is the strain carrying the gene #i whose gene name, locus tag, and accession number are presented in Table S1.

| **Strain** | **Plasmid(s)** | **Source** |
|---|---|---|
| Control (single gene overexpression) | pSR001 | (12) |
| #1 | #1::pSR001 | This study |
| #2 | #2::pSR001 | This study |
| #3 | #3::pSR001 | This study |
| #4 | #4::pSR001 | This study |
| #5 | #5::pSR001 | This study |
| #6 | #6::pSR001 | This study |
| #7 | #7::pSR001 | This study |
| #8 | #8::pSR001 | This study |
| #9 | #9::pSR001 | This study |
| #10 | #10::pSR001 | This study |
| #11 | #11::pSR001 | This study |
| #12 | #12::pSR001 | This study |
| #13 | #13::pSR001 | This study |
| #14 | #14::pSR001 | This study |
| #15 | #15::pSR001 | This study |
| #16 | #16::pSR001 | This study |
| #17 | #17::pSR001 | This study |
| #18 | #18::pSR001 | This study |
| #19 | #19::pSR001 | This study |
| #20 | #20::pSR001 | This study |
| #21 | #21::pSR001 | This study |
| #22 | #22::pSR001 | This study |
| #23 | #23::pSR001 | This study |
| #24 | #24::pSR001 | This study |
| #25 | #25::pSR001 | This study |
| #26 | #26::pSR001 | This study |
| #27 | #27::pSR001 | This study |
| #28 | #28::pSR001 | This study |
| #29 | #29::pSR001 | This study |
| Control (dual-gene overexpression) | pSR001+pSR008 | This study |
| #18+#12 | #18::pSR001+#12::pSR008 | This study |
| #15+#12 | #15::pSR001+#12::pSR008 | This study |
| #18+#16 | #18::pSR001+#16::pSR008 | This study |
| #6+#17 | #6::pSR001+#17::pSR008 | This study |
| #18+#24 | #18::pSR001+#24::pSR008 | This study |
| #1+#1 | #1::pSR001+#1::pSR008 | This study |
| #18+#17 | #18::pSR001+#17::pSR008 | This study |
| #23+#23 | #23::pSR001+#23::pSR008 | This study |
| #16+#17 | #16::pSR001+#17::pSR008 | This study |
| #17+#17 | #17::pSR001+#17::pSR008 | This study |
| #21+#14 | #21::pSR001+#14::pSR008 | This study |

**Supplementary Figure S1.** Gene classification methodology. **Step 1:** For each pairwise set of biological conditions, gene TPM values are floored to a minimum of 5, averaged, and converted into $\log_2$ scale. **Step 2:** Fold change (X and Y) and regulation (Z) scores are calculated between biological conditions. **Step 3:** Genes are classified based on the scores of X, Y and Z into upregulated or increasing for a biological condition, changed regulation, or no change.

#### 1 Average $\text{Log}_2$(TPM) values

| Gene | $\text{BC1}_{\text{early}}$ | $\text{BC1}_{\text{mid}}$ | $\text{BC2}_{\text{early}}$ | $\text{BC2}_{\text{mid}}$ |
|---|---|---|---|---|
| A | # | # | # | # |
| B | # | # | # | # |
| C | # | # | # | # |

| Pairwise Set | Biological condition 1 (BC1) | Biological condition 2 (BC2) |
|---|---|---|
| i | MT 0% IL | WT 0% IL |
| ii | WT 8% IL | WT 0% IL |
| iii | MT 8% IL | MT 0% IL |
| iv | MT 8% IL | WT 8% IL |

#### 2 Calculate X, Y and Z scores

$$\boldsymbol{X} = \frac{\text{BC1}_{early} - \text{BC2}_{early}}{\sqrt{0.25 + (V/N)_{\text{BC1}_{early}} + (V/N)_{\text{BC2}_{early}}}}$$

$$\boldsymbol{Y} = \frac{\text{BC1}_{mid} - \text{BC2}_{mid}}{\sqrt{0.25 + (V/N)_{\text{BC1}_{mid}} + (V/N)_{\text{BC2}_{mid}}}}$$

$$\boldsymbol{Z} = \frac{(\text{BC1}_{mid} - \text{BC1}_{early}) - (\text{BC2}_{mid} - \text{BC2}_{early})}{\sqrt{0.25 + (V/N)_{\text{BC1}_{mid}} + (V/N)_{\text{BC1}_{early}} + (V/N)_{\text{BC2}_{mid}} + (N)_{\text{BC2}_{early}}}}$$

| Variable | Definition |
|---|---|
| X | Fold change at early- exponential sample |
| Y | Fold change at mid-exponential sample |
| Z | Regulation score |
| BC1 | Average $\text{Log}_2$(TPM) of a gene for BC1 |
| BC2 | Average $\text{Log}_2$(TPM) of a gene for BC2 |
| Early | Early-exponential sample |
| Mid | Mid-exponential sample |
| 0.25 | Pseudo variance |
| V | Variance |
| N | Number of replicates |

#### 3 Classify Genes

| Classification | *X* | *Y* | *Z* |
|---|---|---|---|
| BC1 upregulated | X ≥ 1 | Y ≥ 1 | - |
| BC1 increasing | -1 < X < 1 | Y ≥ 1 | Z ≥ 1.5 |
| BC2 upregulated | X ≤ -1 | Y ≤ -1 | - |
| BC2 increasing | -1 < X < 1 | Y ≤ -1 | Z ≤ -1.5 |
| Changed regulation | Is neither upregulated nor increasing | | \| Z \| ≥ 1.5 |
| No change | Is not otherwise classified | | |

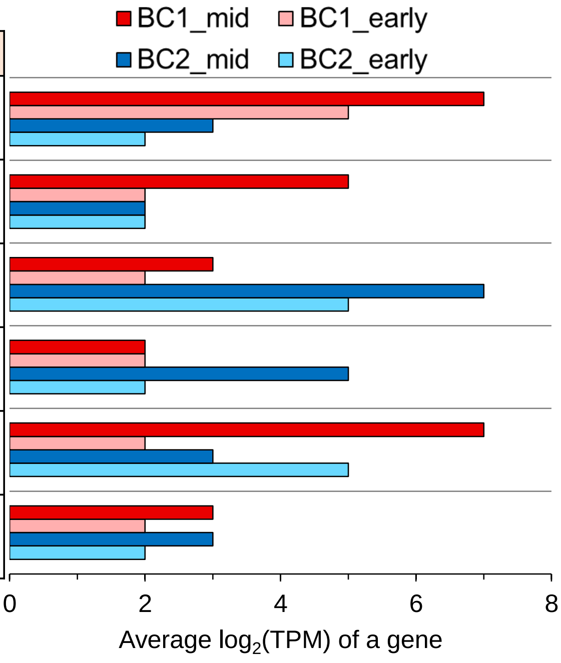
